# Identification of diagnostic KASP markers for selection of crown rust resistance in oats

**DOI:** 10.1101/2024.10.27.620528

**Authors:** Duong T. Nguyen, David Lewis, Eva C. Henningsen, Rohit Mago, Jana Sperschneider, Peter N. Dodds, Melania Figueroa

## Abstract

Oat crown rust, caused by *Puccinia coronata* f. sp. *avenae*, poses a significant threat to global oat production (*Avena sativa* L.). Molecular markers are essential to assist in the integration of multiple resistance genes into a single oat cultivar to achieve genetic resistance durability. Here, we validated previously reported markers for the race-specific resistance genes *Pc39*, *Pc45*/*PcKM*, *Pc54*, and *Pc68*, and developed new KASP markers closely linked to these loci. These markers were subsequently screened across a collection of 150 oat cultivars. Analysis of molecular marker data, pedigree information, and disease resistance profile indicated several oat cultivars likely carrying *Pc39* and *Pc68*. Newly identified carriers of *Pc68* include the cultivars Galileo and Graza80, while Warrego, Kowari, Possum, Forester, Drummond, Hokonui, Glider, and Culgoa were identified as potential carriers of *Pc39*. Discrepancies between the previous postulation of *Pc* gene carriers and our phenotyping and genotyping analysis were also found. For example, the oat line Glider was previously postulated to carry *Pc58* or *Pc59,* but it was positive for the *Pc39*-associated marker and had a similar resistance profile to *Pc39* carriers. These findings underscore the importance of the utilisation of molecular markers in tracking the resistance genes in breeding germplasm.

**Key message:** This study has developed and validated KASP molecular markers associated with crown rust resistance genes *Pc39*, *Pc45*/*PcKM*, *Pc54*, and *Pc68* that could be used for marker-assisted selection in oats.

## Introduction

The fungal disease caused by *Puccinia coronata* f. sp. *avenae* (*Pca*), known as crown rust, is one of the main global threats to oat (*Avena sativa* L.) production (Nazareno et al. 2018; Simons 1985). The infection severity of oat crown rust is influenced by weather conditions, inoculum load, and the genotypes of planted oat cultivars. Epidemics of crown rust can typically reduce oat yields by 10% to 50%; however, entire fields can be lost under optimal environmental conditions (Chambers and Thomas 2020; Strunk et al. 2022). Planting oat cultivars with genetic resistance is considered the most environmentally friendly, and cost-efficient method for rust control (Figueroa et al. 2020, Nazareno et al. 2018).

Race-specific resistance conferred by single-major gene resistance follows a gene-for-gene interaction model (Flor 1971; Dodds 2023) and it has been a common strategy to control crown rust in the field (Nazareno et al. 2018). However, this type of genetic resistance lacks long-term durability as rust populations can rapidly evolve new avirulence (Avr) effector gene variants and avoid plant detection (Figueroa et al. 2020). The evolution of new virulence traits in *Pca* populations and the defeat of resistance has been well-documented in many parts of the world (Henningsen et al. 2024; Klenová-Jiráková et al. 2010; Moreau et al. 2024; Nguyen et al. 2024; van Niekerk et al. 2001) and represents a challenge when breeding for genetic crop protection. In contrast, non-race-specific resistance, also known as partial or adult plant resistance (APR), generally involves broad-spectrum partial resistance that can reduce rust inoculum and therefore in-field disease pressure (Nazareno et al. 2022). Thus, a sound approach to prolong the durability of race-specific genes is the stacking of both types of resistance sources into a single cultivar (Mundt 2018; Periyannan et al. 2017). Marker-assisted selection (MAS) is an effective and high-throughput method to streamline this process (Gnanesh et al. 2013). Nonetheless, the application of MAS in oat breeding for crown rust resistance remains constrained due to the lack of high-quality, and high-throughput markers.

More than 90 plant resistance genes for crown rust have been catalogued in oat (Carson 2017). Among these, a small set of genes (including *Pc39*, *Pc45*, *Pc54*, *Pc68,* and others) have been mapped to chromosome (chr) locations in biparental crosses or by association mapping (Admassu-Yimer et al. 2018, 2022; Bush and Wise 1998; Chen et al. 2006; Gnanesh et al. 2013, 2015; Hoffman et al. 2006; Kebede et al. 2019; Klos et al. 2017; Kulchesiki et al. 2010; McCartney et al. 2011; McNish et al. 2020; Satheeskumar et al. 2011; Sowa and Paczos-Grzeda 2020; Zhao et al. 2020; Wight et al. 2004). However, only a few SNPs linked to these *Pc* genes have been converted into markers that can be readily applied for MAS. The Kompetitive Allele-Specific PCR (KASP) genotyping system for single-nucleotide polymorphisms (SNPs) is one of the most widely adopted technologies that can support MAS in oat breeding (Rahman et al. 2023).

Molecular markers for *Pc39* were mapped to Mrg11 (Sowa and Paczos-Grzeda 2020; Zhao et al. 2020) and several *Pc39-*associated SNP markers were converted into KASP markers and validated in a panel of 74 oat lines and cultivars (Zhao et al. 2020). Among these, the markers *GMI_ES01_c12570_390* and *GMI_ES15_c6153_3929* were proposed to be most useful for MAS. Recently, Wight et al. (2024) used comparative genetic and physical mapping on the oat reference genome OT3098 v2 (PepsiCo, https://wheat.pw.usda.gov/jb?data=/ggds/oat-ot3098v2-pepsico) to assign these KASP markers and several sequence characterised amplified region (SCAR) markers associated with *Pc39* (Sowa and Paczos-Grzęda 2020) to homologous genomic regions on chr4C, chr6A, and chr5D. In another study, Kebede et al. (2019) mapped the gene *Pc45* and confirmed its identity with *PcKM* (Gnanesh et al. 2015). Two KASP markers *I05-0874-KOM16c1* and *GMI_ES02_c16987_268* on Mrg08 (chromosome 2D, OT3098 v2) were reported as the most specific for co-presence/absence of *Pc45*/*PcKM* in a collection of 71 oat lines (Gnanesh et al. 2015; Kebede et al. 2019). The gene *Pc54* was mapped to Mrg02 (chr7D, OT3098 v2) by Admassu-Yimer et al. (2022), with three array-SNPs (*GMI_ES03_c95_413*, *GMI_ES15_c15279_258*, and *GMI_ES22_c2813_554*) reported being closely linked to *Pc54* (Admassu-Yimer et al. 2022). In a genome-wide association study (GWAS), Klos et al. (2017) identified a GBS-derived SNP, *Avgbs_228849*, on Mrg19 (chr3D, OT3098 v2) as significantly associated with *Pc68* resistance to several North American *Pca* isolates.

In this study, we converted SNPs linked to *Pc54* and *Pc68* into KASP markers and used these and the available KASP markers for *Pc39* and *Pc45* to genotype a set of 86 oat crown rust differential lines. This was combined with genome-wide DArTSeq genotypic data from these lines (Nguyen et al. 2023) for association mapping analysis. This analysis validated the previously identified genomic locations for *Pc39*, *Pc45*, *Pc54*, and *Pc68* and allowed the development of new KASP markers specific to these *Pc* genes. A diverse collection of 150 oat lines (Nguyen et al. 2023) was screened using the most specific markers to identify potential sources of genetic resistance. Pedigree information and disease resistance profiles of the oat lines were then compared to further support the putative *Pc* genes present in these oat lines. Overall, the findings from this study provide additional resources for marker-assisted breeding of oat crown rust resistance.

## Materials and methods

### Plant and DNA material

Two oat differential sets from Australia and the USA comprising 86 lines were previously described (Nguyen et al 2023). An oat collection recently compiled by Nguyen et al. (2023) with lines sourced from USDA-ARS (St. Paul, MN, USA), the Australian Grain Genebank (AGG), and an Avena seed stock at CSIRO (**Supplementary file 1**). DNA was extracted from 3-week-old seedlings as described by Ellis et al. (2005) and quality and concentration were assessed using a NanoDrop™ 8000 Spectrophotometer (NanoDrop Technologies Inc., Santa Clara, CA, United States) and PicoGreen fluorescence assay (Singer et al. 1997).

### KASP assays for *Pc39*, *Pc45*/*PcKM*, *Pc54*, and *Pc68*

We adopted the KASP-ready markers for gene *Pc39*, known as *GMI_ES01_c12570_390* and *GMI_ES15_c6153_392* developed by Zhao et al. (2020), and for *Pc45/PcKM*, *GMI_ES02_c16987_268*; *I05-0874-KOM16c1* developed by Gnanesh et al. (2015) and Kebede et al. (2019), respectively. In addition, we designed new KASP markers to the SNPs *GMI_ES15_c15279_258*, *GMI_ES03_c95_413*, *GMI_ES22_c2813_554* that were closely linked with *Pc54* (Admassu-Yimer et al. 2022), and *Avgbs_228849,* that was reported to significantly associate with *Pc68* (Klos et al. 2017). The flanking sequences of these SNPs (extracted from Tinker et al. 2014 and https://oat.triticeaetoolbox.org/) were imported into the Kraken™ software system for marker design using the default parameters (LGC Biosearch Technologies, UK; https://www.biosearchtech.com/). Additional KASP markers *KASP_chr6A_443082340* (*Pc39*), *KASP_chr6A_443082349* (*Pc39*), *KASP_chr6A_443117388* (*Pc39*), *KASP_chr2D_181078165* (*Pc45*), *KASP_chr7D_464499341* (*Pc54*), *KASP_chr3D_466310997* (*Pc68*), and KASP_chr3D_473653096 (*Pc68*) (**Table S1)** were designed based on DArTSeq derived SNPs significantly associated with the genes of interest. KASP assays were performed in the SNPline PCR Genotyping System (LGC, Middlesex, United Kingdom), following the methods described by Shi et al. (2023).

### Association mapping analysis

Genotype association with *Pc* resistance was assessed in the 86 oat differential lines using previously generated genome-wide DArTSeq genotype data for these lines (Nguyen et al., 2023) along with genotypes for eight KASP markers derived from SNPs linked to *Pc39* (Zhao et al., 2020), *Pc45* (Gnanesh et al. 2015; Kebede et al. 2019), *Pc54* (Admassu-Yimer et al. 2022), and *Pc68* (Klos et al. 2017). Each oat line was assigned a phenotype score (either 1 or 2) based on their previous description as carriers (score = 2) or non-carriers (score = 1) of *Pc39*, *Pc45*, *Pc54*, or *Pc68* genes (Carson 2017; Chen et al. 2006; Kebede et al. 2019; Mitchell Fetch et al. 2007; Park 2013). Association analysis was conducted using Trait Analysis by Association, Evolution, and Linkage (TASSEL) software (v5.2.63) (Bradbury et al. 2007). In TASSEL, the population structure and kinship were calculated from the imported marker data using four principal components and Centred Identity by State (IBS) approach, respectively, as inputs for a Mixed Linear Model (MLM) workflow. No compression was performed, and variance components were re-estimated after each marker. Bonferroni correction thresholds were calculated by dividing the *p*-value (α = 0.01) by the number of markers used in association analysis. For each *Pc* gene, the association analysis was limited to SNP markers located on the chromosome(s) to which the genes had previously been mapped (e.g. chr2D for *Pc45*).

### Pedigree investigations

The relationships between oat cultivars were visualised using the Helium pedigree viewing software (Shaw et al. 2014) with pedigree records obtained from the “Pedigrees of Oat Lines” POOL database (Tinker and Deyl 2005) and Fitzsimmons et al. (1983). Pedigrees for each accession were resolved to Parent 1 and Parent 2 to make them compatible with Helium, and the resulting database was then imported into Helium. Together with the pedigrees, previous postulations of resistance genes were imported to Helium for examination in the context of the pedigree.

### Rust infection assays

A set of 16 *Pca* isolates collected from major Australian oat-growing regions in 2022 and 2023 was used (Henningsen et al. 2024; Nguyen et al. 2024). Infection and scoring methods were previously described in Miller et al. (2020). The scores were subsequently converted to a 0-9 numeric scale as described by Miller et al. (2018) and plotted as heatmaps using ComplexHeatmap (v2.14.0) in R (4.2.2) (Gu, 2022).

### Sequence comparison analysis

The physical locations of the markers linked to *Pc* genes of interest were determined by BLAST search of the flanking sequences of the SNPs against the OT3098 v2 reference genome (https://wheat.pw.usda.gov/jb?data=/ggds/oatot3098v2-pepsico) using Geneious Prime® 2022.2.2. The sequences of flanking markers were obtained from NCBI GenBank (https://www.ncbi.nlm.nih.gov/nuccore/) and T3/Oat (https://oat.triticeaetoolbox.org/). Syntenic relationships were established through a Comparative Genomics Platform (CoGe) (https://genomevolution.org). The reference genome OT3098 v2 was imported onto CoGe, and the SynMap2 tool (Haug-Baltzell et al. (2017) was utilised for chromosome alignments and syntenic visualisation https://genomevolution.org/CoGe/SynMap.pl.

## Results

### Identification of new markers for oat crown rust via association mapping analysis

To validate previously reported markers and identify new markers more closely associated with *Pc39, Pc45, Pc54,* and *Pc68,* we conduced an association analysis in 86 lines from the oat crown rust differential sets each carrying defined *Pc* genes. We first scored the genotypes of this set using KASP-ready markers for *Pc39, GMI_ES01_c12570_390* and *GMI_ES15_c6153_392* (Zhao et al. 2020) and *Pc45, GMI_ES02_c16987_268* and *I05-0874-KOM16c1* (Gnanesh et al. 2015), (**Supplementary file 2**). We further designed new KASP markers for *Pc54* and *Pc68* from the closely associated SNPs *GMI_ES15_c15279_258*, *GMI_ES03_c95_413*, *GMI_ES22_c2813_554* (Admassu-Yimer et al. 2022), and *avgbs_228849* (Klos et al. 2017), which successfully detected these SNP genotypes across the differential sets. These marker genotypes were combined with genome-wide DArTSeq SNP genotypes previously generated for these lines (Nguyen et al. 2023) to perform association analysis. Because the chromosome locations of these genes were already known, the association analysis for each gene was restricted to markers on the relevant chromosomes where the genes were previously mapped (**Figure 1** and **Figure S1**).

**Figure 1.**
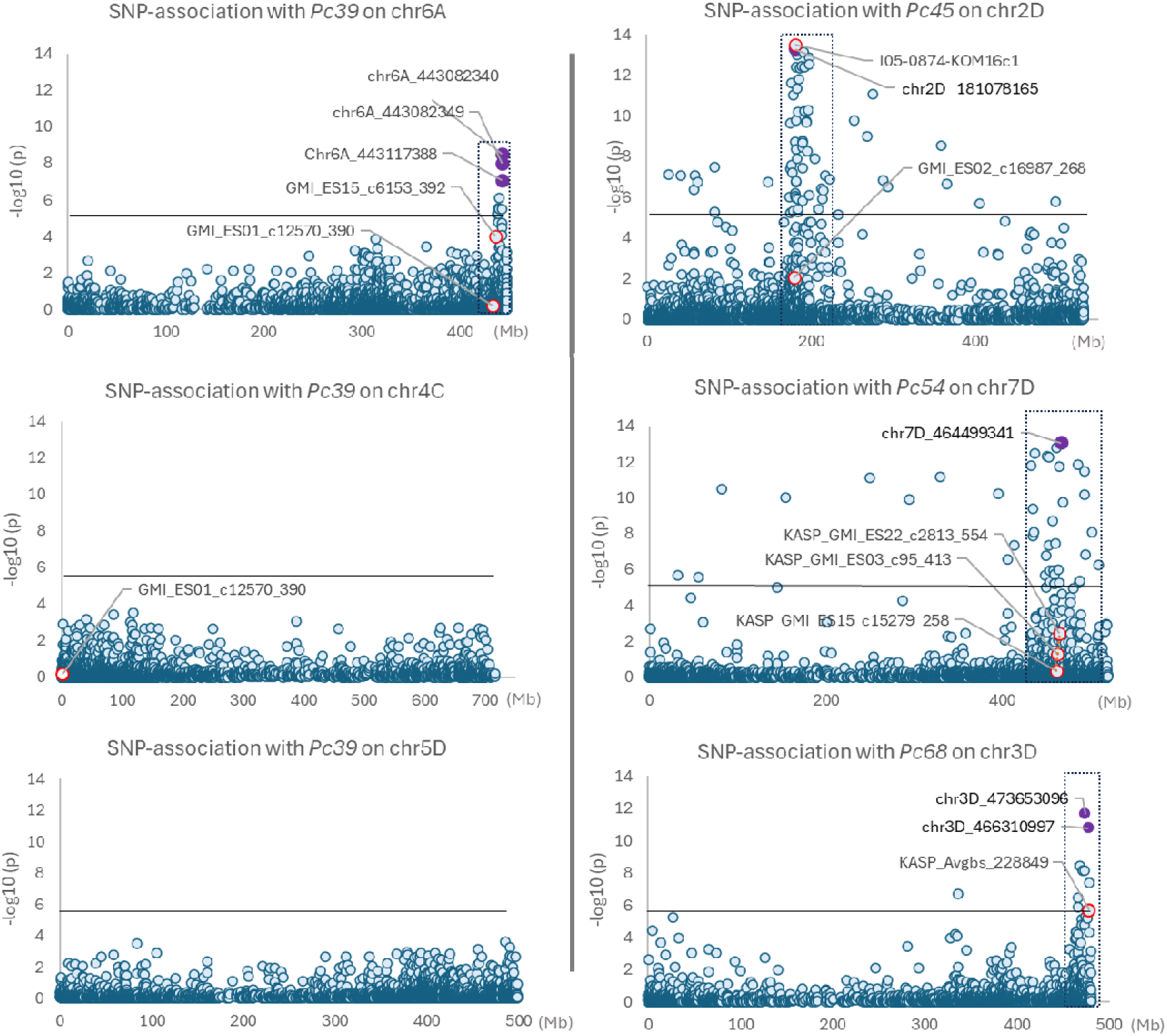
Association analysis for *Pc39*, *Pc45*, *Pc54*, and *Pc68* loci using DArTSeq SNPs (6A = 3641; 4C = 3601; 5D = 2873; 2D = 4604; 7D = 5476; 3D = 3086) and eight published markers in the Differential set (n=86). Red circled dots represent KASP markers for *Pc39*, *Pc45*, *Pc54*, and *Pc68* derived from Zhao et al. (2020), Kebede et al. (2019), Admassu-Yimer et al. (2022) and Klos et al. (2017), respectively. Purple dots are the most significant SNPs selected to convert into KASP markers. Black horizontal lines denote the Bonferroni significance threshold (α= 0.01/total number of markers).

**Figure S1.**
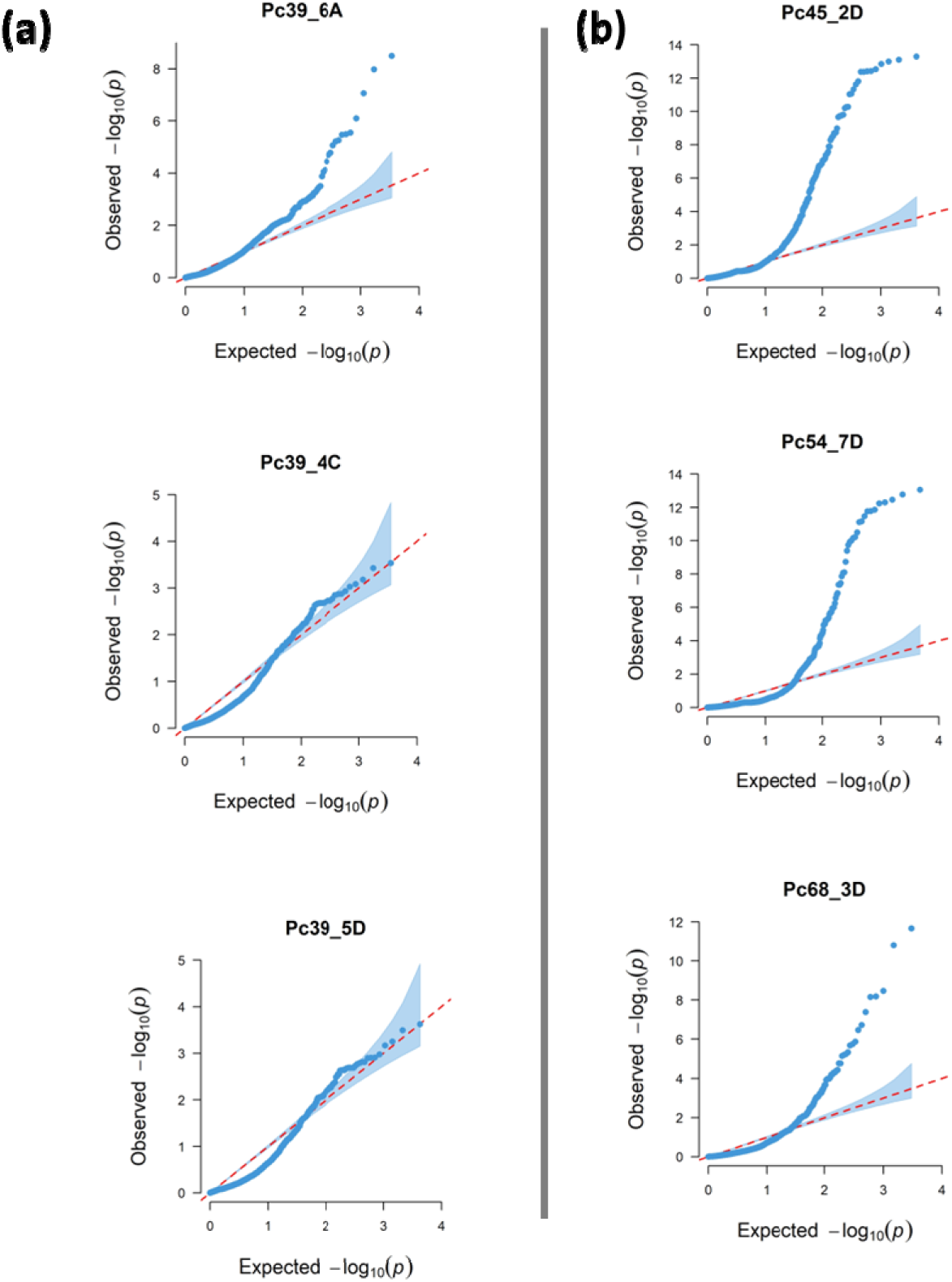
Quantile-quantile plots of SNP association values for carriers of (a) *Pc39* on chromosomes 6A, 4C, and 5D; and (b) *Pc45*, *Pc54*, and *Pc68* on chromosomes 2D, 7D, and 3D respectively. The x-axis and y-axis represent −log10 transformed expected p-values and observed p-values, respectively. The dots indicate −log10(p) of the SNPs and the diagonal line represents the expected values under the null hypothesis for no association.

In a recent report, Wight et al. (2024) assigned the physical positions of several marker sequences that are linked to the *Pc39* locus (Sowa and Paczos-Grzeda 2020; Zhao et al. 2020) to homoeologous regions of chromosomes 4C, 5D, and 6A of reference genome OT3098 v2. Sequence comparison of these chromosomes confirmed that they each contain a relatively small region that is syntenous across all three chromosomes and includes the known marker sequence locations (**Figure S2a**). Therefore, we conducted an association analysis for *Pc39* using DArTSeq markers on chromosomes 4C, 5D, and 6A, along with the KASP markers for this locus. In this analysis, significant association of DArTSeq SNPs with *Pc39* only occurred on chr6A in the region between 436-443 Mb, with no significant associations detected in the homoeologous regions on chromosomes 5D and 4C (**Figure 1**). The markers *GMI_ES01_c12570_390* and *GMI_ES15_c6153_392* (Zhao et al. 2020) both occurred in this region of chr6A, but other DArTSeq SNP markers from this region showed higher association with *Pc68* in this population.

**Figure S2.**
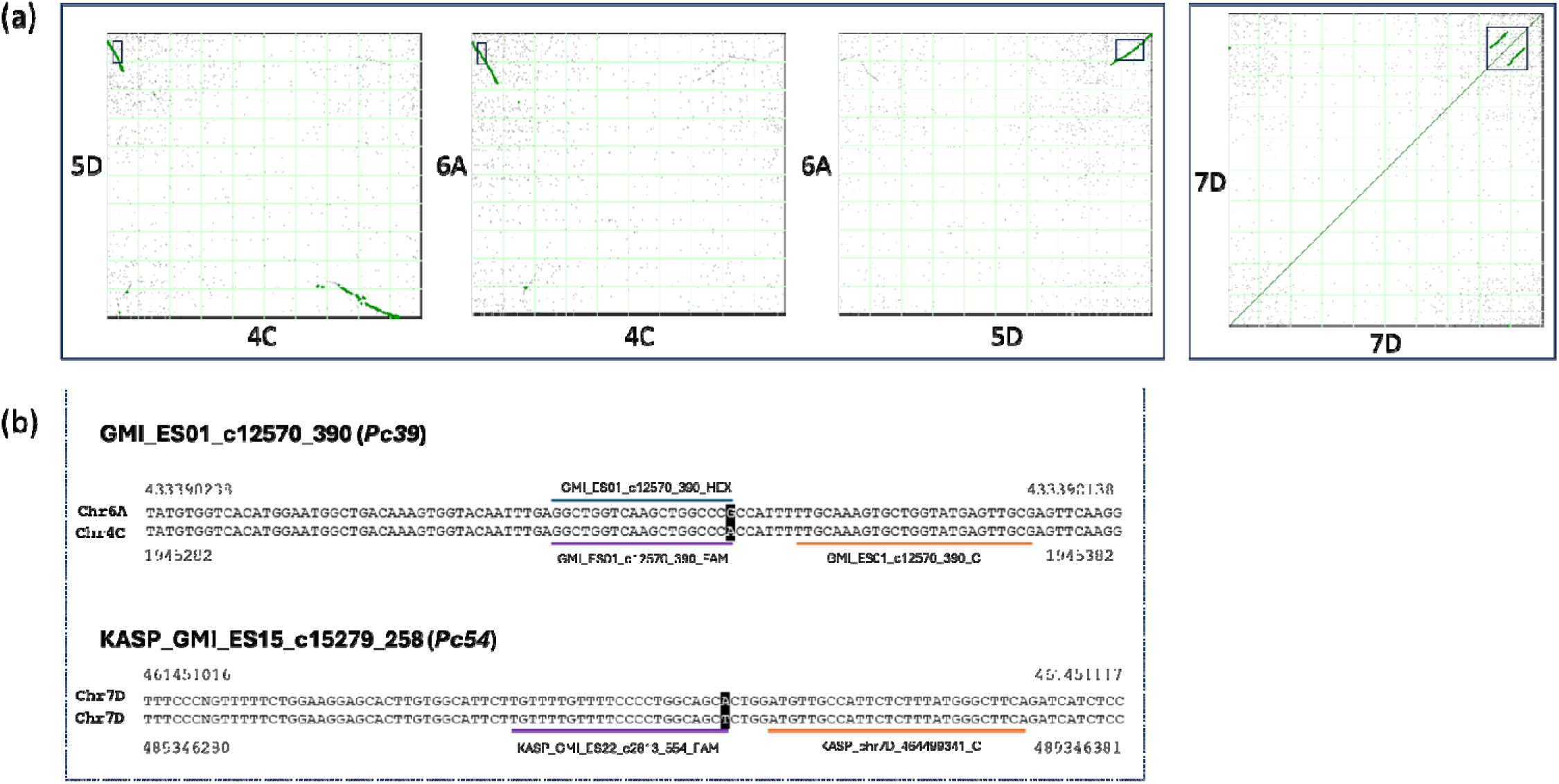
(a) Dot plots comparing chr 6A, 4C, and 5D to each other (left) and 7D to itself (right). The black box indicates homoeologous regions containing *Pc39*-associated markers at ∼1.9 - 27 Mb (chr4C), ∼450 - 490 Mb (chr5D), and ∼400 - 440 Mb (chr6A); and duplicated regions at 441 – 466 Mb and 469 – 494 Mb on chr7D containing *Pc54* associated markers. (b) Comparison of genome sequence BLAST hits for the flanking sequence of markers *GMI_ES01_c12570_390* (*Pc39*) and *GMI_ES15_c15279_258* (*Pc54*) on OT3098 v2 reference genome. Purple and blue underlines indicate KASP allele-specific primer sequences; orange underlines indicate KASP common primer sequences; nucleotides in shades are SNPs. Numbers above/under sequences indicate the physical location.

DArTSeq SNPs associated with Pc68 were located in the 463-478 Mb region of chr3D (**Figure 1**). The marker *KASP_avgbs_228849* for *Pc68* was developed from a GBS-SNP (Klos et al. 2017) and the same SNP was also found as a DArTSeq marker at 470,133,974 bp on chr3D in this study. The genotyping calls for this DArTSeq marker and *KASP_avgbs_228849* were identical, except for a few missing calls in the KASP assay. Several DArTSeq SNP markers showed higher association with *Pc68* than *KASP_avgbs_228849*.

Significantly associated DArTSeq markers for *Pc45* and *Pc54* were found between 180-200 Mb on chr2D and 435-495 Mb on chr7D, respectively (**Figure 1**). These regions are also collocated with the KASP markers corresponding to *Pc45* on chr2D and *Pc54* on chr7D. For *Pc45*, the existing marker *I05-0874-KOM16c1* showed the highest significance scores compared to the DArTSeq SNPs in the association analysis. However, for *Pc54*, several DArTSeq markers showed higher association than the previously described markers.

### KASP marker validation using oat differential sets

From the association analysis, several DArTSeq SNPs most strongly associated with the *Pc* genes were selected and converted into KASP markers (**Figure 1** and **Table S1**). These newly developed KASP markers were genotyped against the differential sets and compared to the results of the previously described markers (**Figure S3)**.

**Figure S3.**
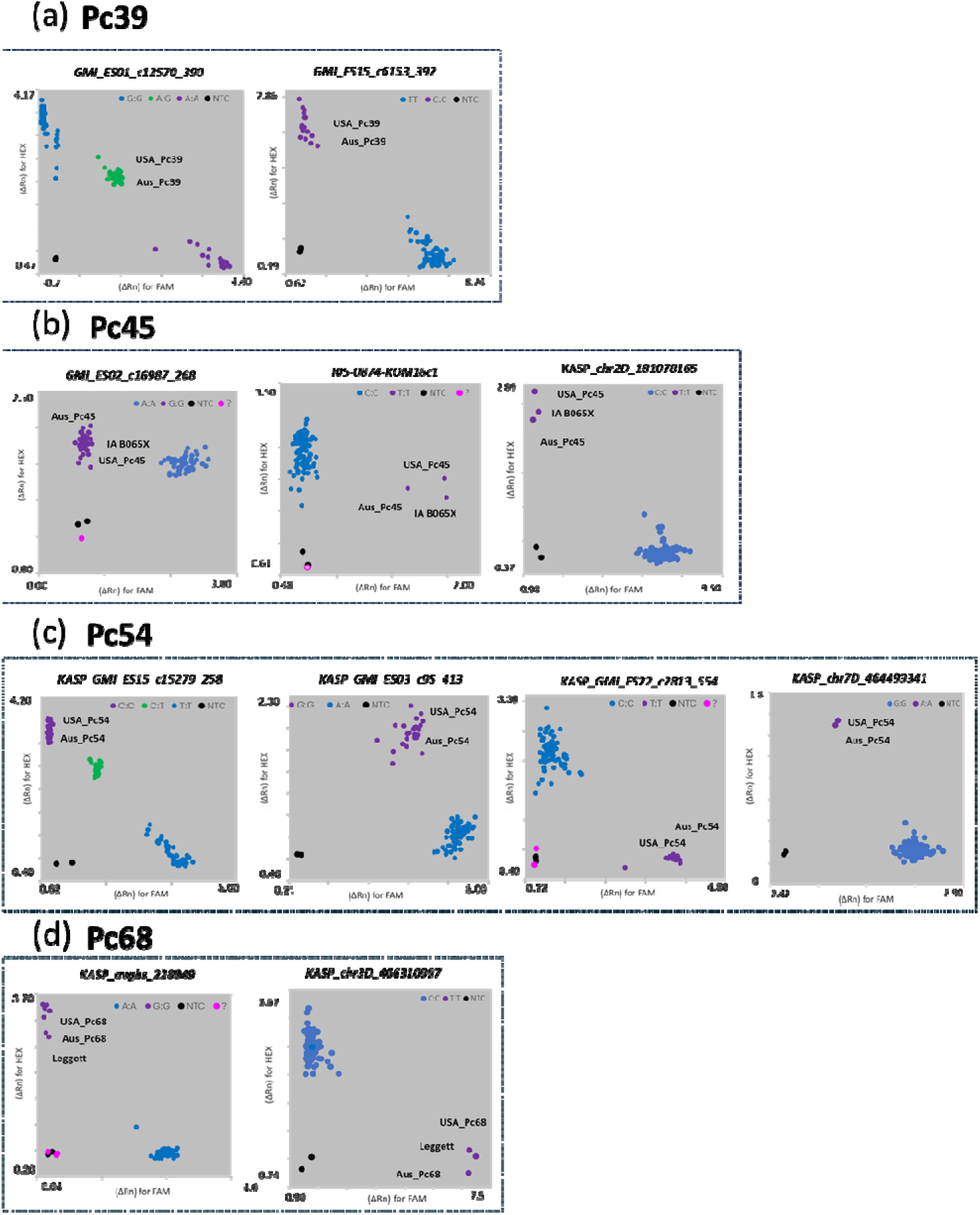
Genotype cluster plots from KASP assays in the differential set (n = 86). (**a**) *Pc39*-linked markers: *GMI_ES15_c6153_392* and *GMI_ES01_c12570_390* (Zhao et al. 2020). (**b**) *Pc45*-linked markers: *I05-0874-KOM16c1* and *GMI_ES02_c16987_268* (Gnanesh et al. 2015), *KASP_chr2D_181078165*. (**c**) *Pc54*-linked markers: *GMI_ES15_c15279_258*, *KASP_GMI_ES03_c95_413*, and *KASP_GMI_ES22_c2813_554* (Admassu-Yimer et al. 2022), *KASP_chr7D_464499341.* (**d**) *Pc68*-linked markers *KASP_avgbs_228849* (Klos et al. 2017), *KASP_chr3D_466310997*. The marker *KASP_chr3D_473653096* for *Pc68* (not shown) displayed a similar pattern as *KASP_chr3D_466310997*. Purple and blue dots show homozygous genotypes for resistant and susceptible alleles; green dots show heterozygous genotypes; black dots represent non-template control; pink dots show ambiguous signals. X- and Y-axes show relative amplification units (ΔRn) for FAM and HEX signals.

#### Pc39

The KASP assay of the marker *GMI_ES01_c12570_390* displayed three distinct genotypes (*AA*/*AG*/*GG*) among the differential lines (**Figure S3a** and **Supplementary file 2**). The presence of the apparently heterozygous genotype (*AG*) in the Pc39 differential lines and 33 other lines might be attributed to the amplification of similar sequences on chromosomes 4C and 6A that differ by this SNP within individual oat lines, as is the case in the OT3098 v2 reference (**Figure S2b**). The occurrence of homoeo-SNPs is a common phenomenon in polyploid species, often resulting in a reduced validation efficiency of the KASP assay (Mammadov et al. 2012). The marker *GMI_ES01_c12570_390* is, therefore, considered non-effective for *Pc39* discrimination in the KASP assay.

On the other hand, the *GMI_ES15_c6153_392* KASP marker was more specific than *GMI_ES01_c12570_390*, detecting 17 positive differential lines compared to 52 (**Table 1** and **Supplementary file 2**). Of the *GMI_ES15_c6153_392* positive lines, two are the defined *Pc39* differential lines from the USA and AUS sets, and two others, Cleanleaf and Leggett, were postulated to carry *Pc39* based on phenotyping (Mitchell Fetch et al. 2007; Park 2013). However, this marker was not positive in Barcoo, which was previously postulated to carry *Pc39*, *Pc61*, and *PcBett* (Park 2013). The other positive lines either carry different resistance genes (e.g. *Pc45*, *Pc58*, *Pc60*, *Pc61*, *Pc70*, *Pc91*, *Pc96*) or unknown genes, such as Culgoa (*PcCulgoa*), Genie, and Warrego (both postulated to have *Pc61* + unknown gene(s) – by Park et al. 2009) (**Supplementary file 2**). A search on the ‘Pedigree of Oat Lines’ POOL database (https://triticeaetoolbox.org/POOL/index_db.php), revealed pedigree connections between Genie, Warrego, HiFi, and Pc91 (ND894904) with other Pc39 carriers (**Figure 2**). The three KASP assays designed from

**Figure 2.**
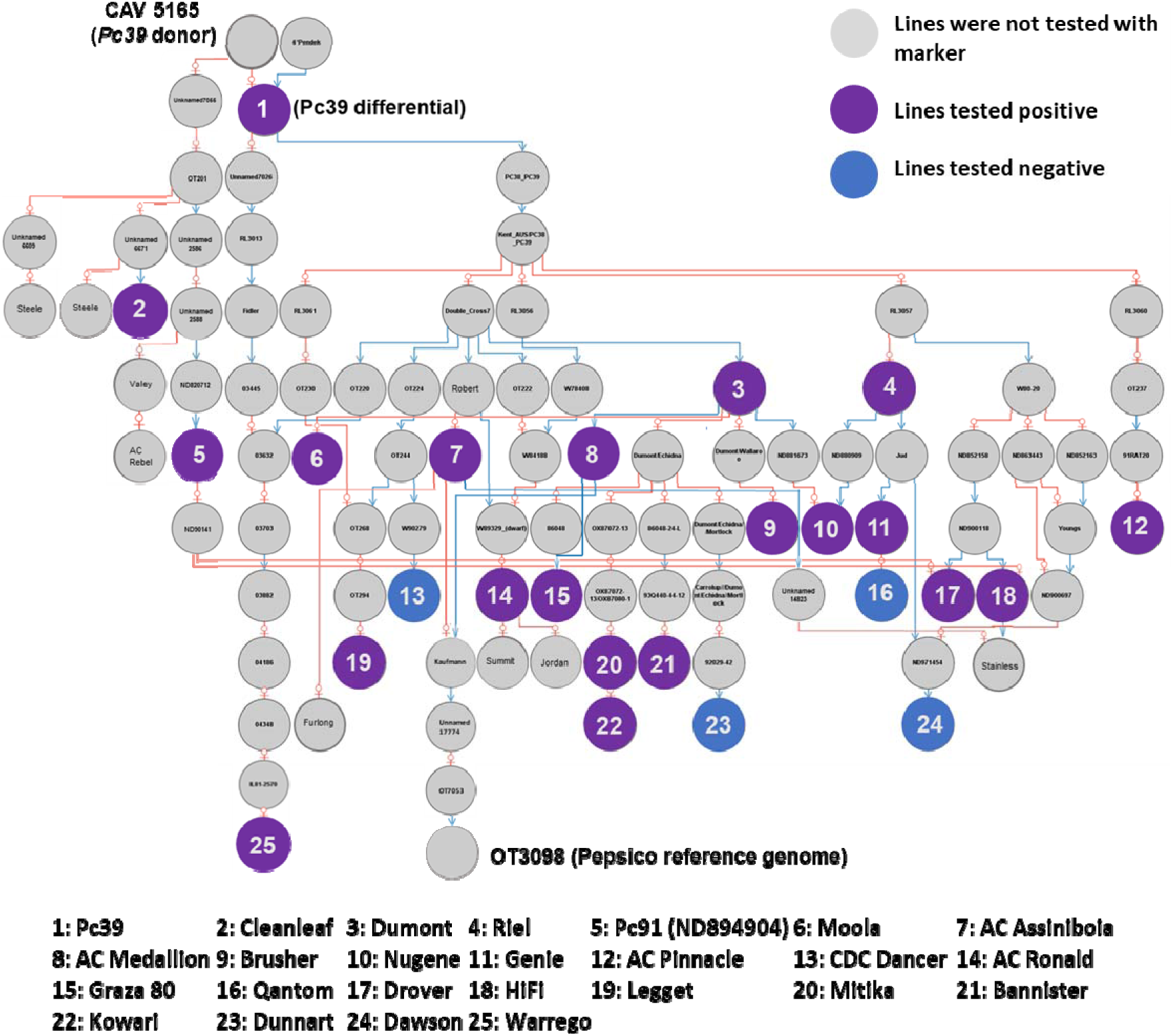
Pedigree of oat lines related to the *Pc39* donor (CAV 5165), modified from a Helium output (Shaw et al. 2014). Pedigrees were obtained from the “Pedigrees of Oat Lines” POOL database (https://triticeaetoolbox.org/POOL; Tinker and Deyl, 2005) and Fitzsimmons et al. (1983). Red lines indicate maternal parent and blue indicates paternal parent. Coloured circles represent lines that tested positive (purple), negative (blue), and were not tested (grey) with the *Pc39* marker *GMI_ES15_c6153_392*.

**Table 1:** KASP marker validation results in the differential sets (n=86). Marker names starting with “KASP” were designed in this study. The “/” separates the locations with equally highest confidence from blast search in reference genome OT3098 V2.

| Pc gene | Reference | Marker | Physical location (OT3098 V2) | Number of lines positive for resistance allele in differential set (n=86) |
| --- | --- | --- | --- | --- |
| Pc39 | Zhao et al. (2020) | *GMI_ES01_c12570_390* | chr6A: 433390147-433390201/<br>chr4C: 433390138-433390238 | 52* |
|  |  | *GMI_ES15_c6153_392* | chr6A: 436047504-436047387 | 17 |
| Pc45/PcKm | Kebede et al. (2019) | *GMI_ES02_c16987_268* | chr2D: 179682216-179682335 | 43 |
|  |  | *I05-0874-KOM16c1* | chr2D: 181118507-181118369 | 3 |
|  | This study | *KASP_chr2D_181078165* | chr2D: 181078104-181078224 | 3 |
| Pc54 | Admassu-Yimer et al. (2022) and this study | *KASP_GMI_ES15_c15279_258* | chr7D: 461451176-461450972/<br>chr7D: 489346440-489346236 | 44* |
|  |  | *KASP_GMI_ES03_c95_413* | chr7D: 462940037-462939953/<br>chr7D: 490417282-490417198 | 24 |
|  |  | *KASP_GMI_ES22_c2813_554* | chr7D: 465112841-465112723/<br>chr7D: 494022361-494022243 | 12 |
|  | This study | *KASP_chr7D_464499341* | chr7D: 464499281-464499400/<br>chr7D: 493116025-493116144 | 2 |
| Pc68 | Klos et al. (2017) | *KASP_Avgbs_228849* | chr3D: 470133926-470134034 | 8 |
|  | This study | *KASP_chr3D_466310997* | chr3D: 466310942-466311050 | 3 |
|  |  | *KASP_chr3D_473653096* | chr3D: 473653029-473653163 | 3 |

\* Indicates the presence of heterozygosity in some of the lines that could be due to duplication of the marker sequence in homoeo-alleles or locally.

Three DArTSeq SNPs (*chr6A_443082340*, *chr6A_443082349, chr6A_443117388*) showed considerably higher association with *Pc39* than the existing KASP markers. However, new KASP markers generated from these SNPs detected only the resistance-associated SNP genotype in all tested lines (Data not shown), which is not consistent with the DArTSeq genotyping data. The challenge for the development of valid markers for *Pc39* may arise from the homoeologous nature of the *Pc39* mapping region, which could have led to primers not being specific to a single chromosome or sequencing assembly error in the OT3098 v2 reference genome.

#### Pc45

A comparison of the genotyping results for *KASP_chr2D_181078165* (developed from DArTSeq SNP in this study) with the markers *I05-0874-KOM16c1* and *GMI_ES02_c16987_268* (Gnanesh et al. 2015; Kebede et al. 2019) on the differential set showed that *KASP_chr2D_181078165* and *I05-0874-KOM16c1* were equally specific to *Pc45*/*PcKM* (**Table 1** and **Supplementary file 2**), as they were both only positive for the two Pc45 differential lines (USA and AUS), plus one additional oat line, IA B605X, which is the *PcKM* donor (Gnanesh et al. 2015; McMullen et al. 2005). In contrast, *GMI_ES02_c16987_268* was not specific for *Pc45*/*PcKM*, as the resistance allele also appeared in 34 other oat lines, that were postulated to possess other sources of resistance (**Supplementary file 2**).

#### Pc54

The new marker *KASP_chr7D_464499341* demonstrated higher specificity for Pc54 than the previously published markers *GMI_ES03_c95_413*, *GMI_ES15_c15279_258*, and *GMI_ES22_c2813_554* (Admassu-Yimer et al. 2022) (**Table 1** and **Figure S3c**) as it was only positive for the two Pc54 differential lines in this set. In contrast, marker *KASP_GMI_ES22_c2813_554* was positive for 12 lines (**Table 1**), including the Pc54 differential lines and ten other lines (**Supplementary file 2**), while the resistance allele of *KASP_GMI_ES03_c95_413 KASP_GMI_ES15_c15279_258*, was detected in 24 and 44 oat lines, respectively, in addition to the Pc54 differential lines. All *Pc54* marker sequences were found at two identical locations near the distal end of chr7D, approximately 28 Mbp apart (**Table S1**). Chromosome alignment indicated a large tandem duplication in this region of chr7D (**Figure S2a**). Moreover, the marker *KASP_GMI_ES15_c15279_258* exhibited heterozygosity in some lines (**Figure 3c**). This is likely a consequence of polymorphism between the duplicated regions on chr7D, as is the case in the OT3098 reference genome (**Figure S2b**).

**Figure 3.**
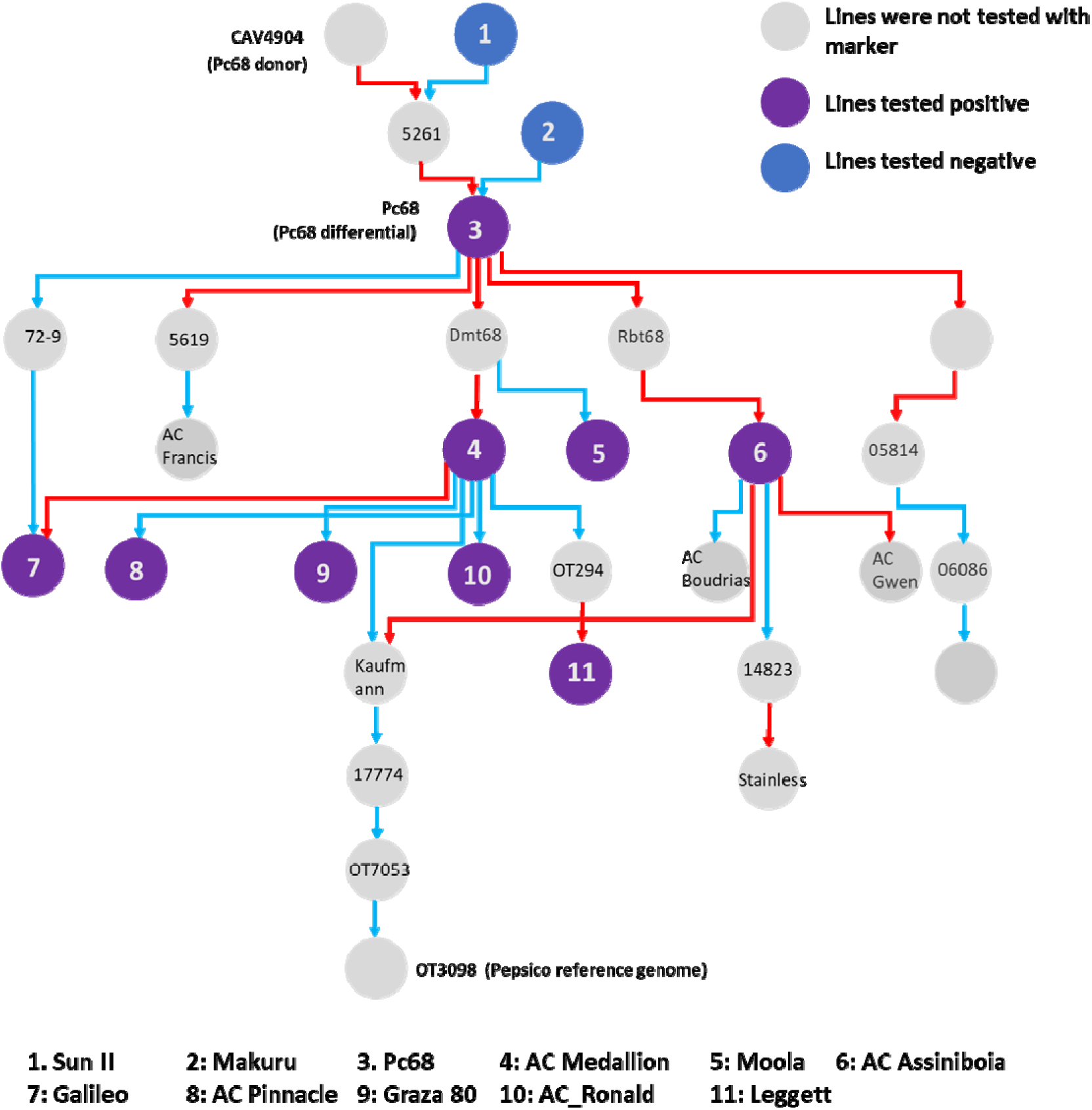
Pedigree of *Pc68* differential, modified from a Helium output (Shaw et al. 2014). Pedigrees were obtained from the “Pedigrees of Oat Lines” POOL database (https://triticeaetoolbox.org/POOL; Tinker and Deyl, 2005) and Fitzsimmons et al. (1983). Red lines indicate maternal parent and blue indicates paternal parent. Coloured circles represent lines that tested positive (purple), negative (blue), and were not tested (grey) with the *Pc68* marker *KASP_chr3D_466310997* and *KASP_chr3D_473653096*.

#### Pc68

The two new *Pc68* markers, *KASP_chr3D_473653096* and *KASP_chr3D_466310997*, are highly specific to known *Pc68* carriers, detecting only the two Pc68 differential lines and the oat line Leggett postulated to carry *Pc39*, *Pc68*, and *Pc94* (Mitchell Fetch et al. 2007). In contrast, the published marker *KASP_avgbs_228849* (Klos et al. 2017) was positive for these three lines as well as five additional lines: Amagalon (*Pc91*), Bondvic, H548, Trispernia (*Pc6D*), and H441 (*Pc53*), none of which have been postulated to carry *Pc68*, and no pedigree connection to *Pc68* carriers was found for these lines (https://triticeaetoolbox.org/POOL/index_db.php) (**Table 1** and **Supplementary file 2**).

### Implementation of diagnostic KASP markers for oat crown rust resistance genes in screening oat collections

After validation of KASP markers in the differential sets, the most specific markers for *Pc39* (*GMI_ES15_c6153_392*), *Pc45* (*I05-0874-KOM16c1*, *KASP_2D_181078165*), *Pc54* (*KASP_chr7D_464499341*) and *Pc68* (*KASP_chr3D_466310997, KASP_chr3D_473653096*) were used to screen a broader collection of 150 oat cultivars to identify potential carriers of these *Pc* genes.

The marker *GMI_ES15_c6153_392* (Zhao et al. 2020) identified 23 oat lines positive for the *Pc39* resistance allele in the oat collection. Among these 16 have pedigree connections with known *Pc39* carriers (**Figure 2** and **Supplementary file 2**) and 10 were previously postulated to carry *Pc39*, such as Dumont, Moola (AC Medallion), AC Assiniboia, Riel, AC Ronald, AC Pinnacle, Leggett, Mitika, Brusher, and Bannister (Mitchell Fetch et al. 2007; Sowa and Paczos-Grzęda 2020; Park et al. 2009; Park et al. 2023). A rust phenotyping experiment using 16 *Pca* isolates (Henningsen et al. 2024; Nguyen et al. 2024) was conducted for 13 lines that were positive for the *Pc39* marker but not previously postulated to carry *Pc39* (**Figure S4a**). Four oat lines, Kowari, Forester, Drummond, and Hokonui, displayed similar resistance profiles to the *Pc39* differential. Five other positive lines (Drover, Warrego, Culgoe, Glider and Possum) were resistant to all *Pc39* avirulent isolates plus one or more additional isolates. Of these, Drover and Warrego both have pedigree connections with the Pc39 differential. This suggests that *Pc39* might be part of resistance mechanisms in these lines. Drover is also postulated to carry Pc91 (Nguyen et al., 2024). Notably, Glider and Warrego were previously postulated to carry *Pc58*/*Pc59* and *Pc61*+, respectively but exhibited quite different resistance profiles than the Pc58, Pc59, and Pc61 differential lines, indicating that *Pc39* and/or other genes, rather than the previously postulated genes may contribute to the resistance in these lines. Culgoa and Possum were previously postulated to contain unknown Pc genes. Lastly, four oat lines Nugene (*PcNugene)*, Genie (*Pc61+*), Graza 80, and Cypress showed similar resistance profiles to *Pc39* but were susceptible to at least one Pca isolate that was avirulent to Pc39. This variation raises uncertainty as to whether these lines carry the *Pc39* resistance gene, despite all testing positive for the *Pc39* marker and three (Nugene, Genie, and Graza 80) sharing pedigree connections with known *Pc39* carriers (**Figure 2**).

**Figure S4.**
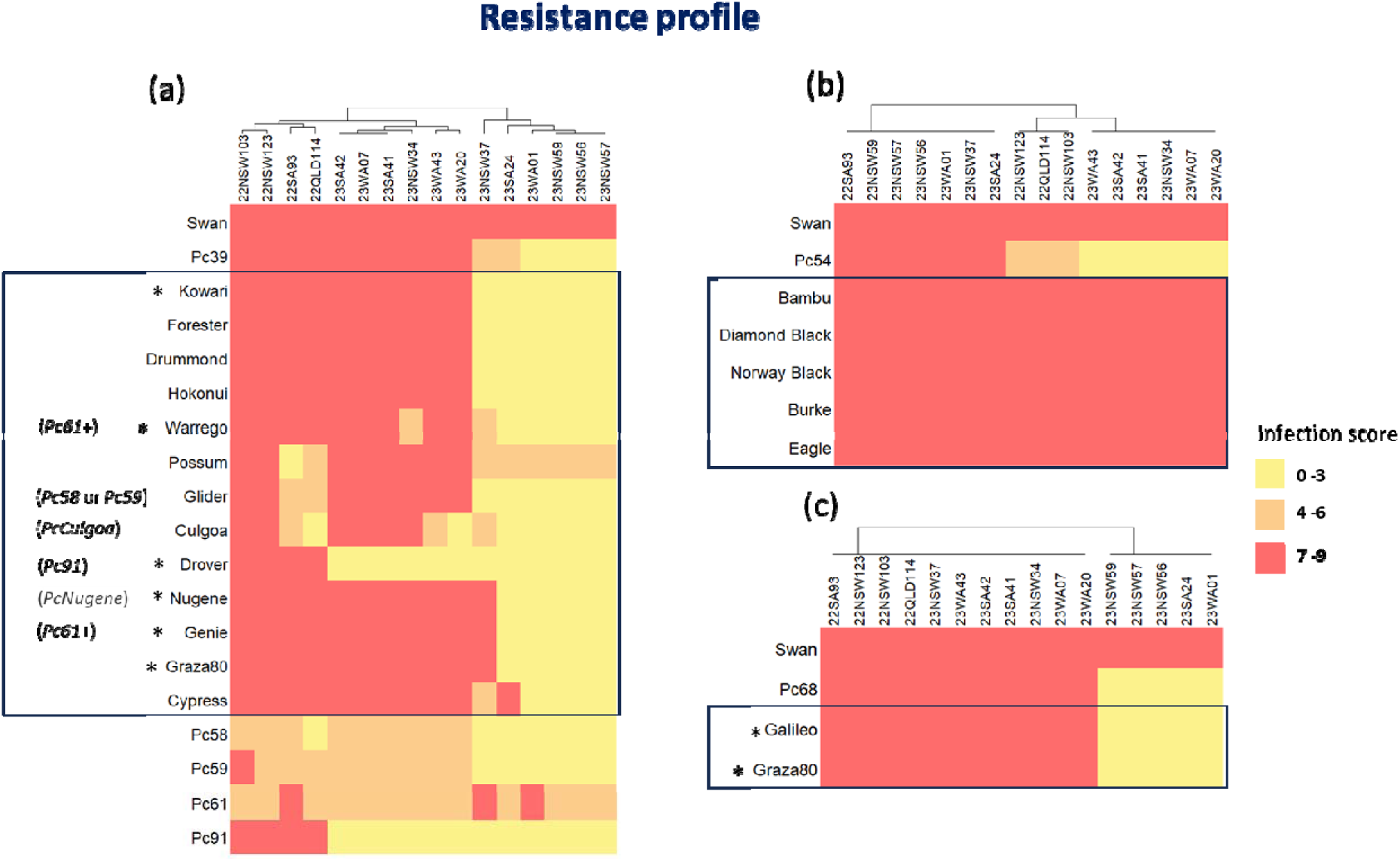
Heatmap comparing resistance profiles against 16 *Pca* isolates of lines that tested positive for (a) *Pc39*, (b) *Pc54*, and (c) *Pc68* markers, but never postulated as carriers (in black boxes). Columns are ordered by hierarchical clustering. * Indicate lines that shared pedigree connections with *Pc* genes of interest. High infection scores indicating high virulence (susceptibility) are shown in red, and lower infection scores indicating avirulence (resistance) are shown in yellow and orange. In single brackets are previously postulated genes in rust reports based on phenotype (Park et al. 2009).

The *Pc45-*associated marker *I05-0874-KOM16c1* derived from Gnanesh et al. (2015) and the marker *KASP_2D_*181078165, derived from the association analysis were not positive for any line in the oat collection (**Supplementary file 2**). This implied the absence of *Pc45* resistance alleles in these oat cultivars.

The *Pc54*-associated marker *KASP_chr7D_46449934*, identified five lines, Bambu, Diamond Black, Norway Black, Burke, and Eagle as potential carriers of *Pc54*. However, no pedigree connections were seen between these lines and the *Pc54* donor (CAV1832). In addition, these lines also exhibited different resistance profiles to the Pc54 differential line (**Figure S4b**), suggesting that they do not carry this resistance gene.

The two *Pc68*-associated markers *KASP_chr3D_466310997* and *KASP_chr3D_473653096* showed identical genotype profiles across the set of oat lines and both identified seven positive lines (**Supplementary file 2**). Five of these, Leggett, AC Ronald, AC Pinnacle, Moola (AC Medalion), and AC Assiniboia, were previously reported as *Pc68* carriers (Chen et al. 2006), while Galileo and Graza 80 are likely also carriers based on their pedigree connection to other *Pc68* carriers (**Figure 3**) and similar resistance profile to the *Pc68* differential line (**Figure S4c**). The oat lines CDC Dancer, CDC Boyer, Starter, Makuru, Sun II, Swan, Riel, and Ogle were previously reported as non-carriers of *Pc68* (Chen et al. 2006; Kulcheski et al. 2010, Wong et al. 1983; Satheeskumar et al. 2011), and they were also negative for markers *KASP_chr3D_466310997* and *KASP_chr3D_473653096*.

## Discussion

Molecular markers diagnostic for specific genes allow breeders to incorporate multiple distinct sources of resistance into the same variety. To date, only a limited number of KASP markers associated with oat crown rust resistance have been validated for use in breeding programs. The present study leveraged gene mapping information (Admassu-Yimer et al. 2022; Gnanesh et al. 2015; Kebede et al. 2019; Klos et al. 2017; Zhao et al. 2020) and DArTSeq genotypic data (Nguyen et al. 2023) to validate markers and genetic locations for *Pc39*, *Pc45*, *Pc54*, and *Pc68* and develop improved KASP markers for future use in the breeding of oat crown rust resistance. The markers most specific for the presence of *Pc39*, *Pc45*, *Pc54*, and *P68* in the differential sets were found to be: *GMI_ES15_c6153_392* (Zhao et al. 2020)) for Pc39; I05-0874-KOM16c1 (Gnanesh et al. 2015) and KASP_chr2D_181078165 (this study) for Pc45/PcKM; KASP_chr7D_464499341 (this study) for Pc54; and KASP_chr3D_473653096 and KASP_chr3D_466310997 (this study) for Pc68 (**Table 1**).

Among the lines positive for the *Pc39* marker *GMI_ES15_c6153_392*, differential lines Pc61 (USA) and Pc70 (USA) were closely related to Pc39 differentials (Aus and USA) on the phylogenetic tree (Nguyen et al. 2023). Moreover, Pc70 (USA) was proposed to be a duplicate of the Pc39 differential as they had the same resistance profile (Henningsen et al. 2024). The Pc91 differential lines and other *Pc91* carriers (HiFi and Drover) have pedigree connections to the Pc39 differential and all tested positive for the *Pc39* marker. This aligns with a recent GWAS analysis of an oat crown rust pathogen population (Hewitt et al. 2023), which identified several genomic regions in *Pca* associated with virulence on *Pc91*, one of which is also associated with virulence on *Pc39*. A possible explanation for this is that the Pc91 differential line and its derivatives (HiFi and Drover) may carry multiple resistance genes including *Pc39* within their genetic background. On the other hand, *Pc39, Pc55*, and *Pc71* were proposed to be different alleles of the same gene based on their similar resistance profiles to *Pca* populations in the USA and Canada (Chong and Seaman 1989; Miller et al. 2020; Leonard et al. 2005). Additionally, Klos et al. (2017) mapped *Pc71* to the same homoeologous regions as *Pc39* (Chr4C, chr5D, and chr6A, OT3098 v2). In our study, Pc55 and Pc71 differential lines were both negative for the *Pc39* marker *GMI_ES15_c6153_392* (**Supplementary file 2**). This indicates that the marker is specific for *Pc39* but not *Pc55* and *Pc71*. The oat cultivar Durack was previously thought to carry *Pc39* (Park et al. 2023), but it tested negative for the *Pc39* marker *GMI_ES15_c6153_392* (**Supplementary file 1**) and showed no pedigree connection to known *Pc39* carriers. Similarly, the cultivar Barcoo, postulated to carry *Pc39*, *Pc61*, and *PcBett* (Park 2013), also tested negative for *Pc39* markers (**Supplementary file 2**) and lacks pedigree connections to Pc39 carriers. Furthermore, Barcoo displayed resistance to all tested *Pca* isolates from Australia in 2022 and 2023 (Henningsen et al. 2024, Nguyen et al. 2024), unlike the differential lines Pc39, Pc61, and Bettong (*PcBett*), which were broadly susceptible to *Pca* isolates in 2023. This suggests that Barcoo’s resistance may be attributed to other or additional genes.

Previously, the KASP marker *I05-0874-KOM16c1,* located at 181,118,479 Mb on chromosome 2D (OT3098 v2), was found to be closely linked with *Pc45*/*PcKM* in five mapping populations and was reported to have the best predictive power for the presence/absence of *Pc45* and *PcKM* in a panel of 71 diverse oat lines (Kebede et al. 2019). In this study, *I05-0874-KOM16c1* and the newly developed marker for *Pc45*/*PcKM*, *KASP_chr2D_181078165* derived from association analysis (approximately 40Kb distant from *I05-0874-KOM16c1)* in the differential sets, showed an equivalently high diagnostic level, detecting only the Pc45/PcKM differentials within this set (**Supplementary file 2**). Both markers detected only susceptibility-assoicated alleles when screening an oat collection of 150 lines, suggesting the absence of *Pc45*/*PcKM* in these stocks. Admassu-Yimer et al. (2018) demonstrated a close linkage between *Pc45* and *Pc53* within 1cM on Mrg08 (chr2D), suggesting they are either tightly linked or allelic. In this study, the Pc53 differential line was negative for all *Pc45*/*PcKM* markers (**Supplementary file 2**), indicating that these markers differentiate between these linked resistance genes.

The marker *KASP_chr7D_464499341* was also highly diagnostic for *Pc54,* detecting only the two designated Pc54 lines within the differential set (**Table 1**). This and other markers linked to *Pc54* (Admassu-Yimer et al. 2022) were identified within tandem duplicated regions on the distal end of chr7D - OT3098 v2 (**Figure S4**). This duplication was also observed by Wight et al. (2024), who suggested it to contain the rust resistance gene *Pc38* (mapped by Wight et al. 2004) through comparative mapping. Additionally, *Pc54* has been previously suggested to be allelic or closely linked to *Pc35* (Simon 1978). In our current investigation, the differential lines Pc38 and Pc35 were negative for three and four of the *Pc54* markers, respectively (**Supplementary file 2**). This outcome indicates that the markers effectively differentiate *Pc54* from the two genes *Pc35* and *Pc38.* Screening of the Oat collection with *KASP_chr7D_464499341* revealed the resistance allele of *Pc54* to be rare, only appearing in five oat lines. However, these showed no pedigree connection to Pc54 and quite different resistance profiles suggesting that these lines are not the carriers of *Pc54*, and there might have been recombination between the marker and the causative gene in these lines. Nevertheless, this marker is suitable for selection of *Pc54* in most genetic backgrounds tested.

The *Pc68* marker *KASP_avgbs_228849*, derived from the GBS-SNP *avgbs_228849* (Klos et al. 2017), along with two newly developed markers, *KASP_chr3D_466310997* and *KASP_chr3D_473653096* from DArTSeq SNPs, are located within the *Pc68* associated region previously identified by Kulcheski et al. (2010), Chen et al. (2006), and Satheeskumar et al. (2011). Of these, *KASP_chr3D_466310997* and *KASP_chr3D_473653096,* accurately distinguished known *Pc68* carriers from non-carriers in the oat collection (**Supplementary file 2**). In contrast, according to Chen et al. (2006) their STS and SNP markers incorrectly identified CDC Dancer and CDC Boyer as *Pc68* carriers, despite these lines not possessing the trait.

Notably, the lines Graza 80 and Galileo, which had not been postulated to have Pc68, were also positive for the two new *Pc68* markers and displayed similar resistance profiles to the Pc68 carrier. Both lines are descendants of AC Medallion (a known *Pc68* carrier), suggesting that these two likely also carry *Pc68*. The two newly developed KASP markers for *Pc68* are deemed to be diagnostic for *Pc68* selection.

In conclusion, our association analysis within oat crown rust differential sets confirmed the physical locations of *Pc39*, *Pc45*/*PcKM*, *Pc54*, and *Pc68*. KASP markers linked to the resistance genes, developed from this and previous studies, were validated as highly diagnostic for the designated Pc genes and identified several potential carriers of *Pc39* and *Pc68* in a diverse oat collection. The outcomes of this study offer valuable tools for MAS in breeding programs to facilitate integration of these resistance loci into oat varieties and generation of lines with combinations of crown rust resistance genes.

## Supporting information

Supplementary file 1. List of plant materials

Supplementary file 2. KASP genotyping results

Table S1. KASP marker sequences

## Statements & Declarations

### Author contributions

This study was planned and designed by MF and PND. ECH and DL manage collections of rust isolates, virulence assessments. DTN performed genotyping and genotypic analysis and generated figures. RM assisted with marker design. JS supported computational analysis. DTN, MF, and PND wrote the first draft of the manuscript. All authors contributed to the interpretation of results, reviewed the manuscript, and approved the final version.

## Acknowledgments

We acknowledge the Australian Grains Genebank, Shahryar F. Kianian at the US Department of Agriculture-Agricultural Research Service, and the GRDC-funded oat phenology project team (grant CSP2007) for contributing oat germplasm. We thank Ben Trevaskis (CSIRO) for helping with pedigree information of several oat cultivars and Liza Apps for technical support at the CSIRO Black Mountain Laboratories Quarantine facility. We also thank Scott Boden at the University of Adelaide for providing DNA material for a subset of oat lines.

## Literature Cited

Admassu-Yimer B, Bonman JM, Esvelt Klos K (2018.) Mapping of crown rust resistance gene *Pc53* in oat (*Avena sativa*). PLOS ONE. 10.1371/journal.pone.0209105

Admassu-Yimer B, Klos KE, Griffiths I et al (2022). Mapping of crown rust (*Puccinia coronata* f. sp. *avenae*) resistance gene *Pc54* and a novel quantitative trait locus effective against powdery mildew (*Blumeria graminis*f. sp.*avenae*) in the oat (*Avena sativa*) line Pc54. Phytopathology 112:1316–1322. 10.1094/PHYTO-10-21-0445-R

Bush AL, Wise RP (1998). High-resolution mapping adjacent to the *Pc71* crown-rust resistance locus in hexaploid oat. Molecular Breeding 4: 13–21. 10.1023/A:1009652222382

Carson M (2017, October 4). Oat crown rust. Cereal Disease Lab, Agricultural Research Service, USDA. https://www.ars.usda.gov/midwest-area/stpaul/cereal-disease-lab/docs/cereal-rusts/oat-crown-rust/

Chambers K, Thomas G (2020). Plant diseases impacting oaten hay production in Australia – a review. Department of Primary Industries and Regional Development (DPIRD). https://www.agric.wa.gov.au/oats/2020-plant-diseases-impacting-oaten-hay-production-australia

Chen G, Chong, J, Gray M, Prashar S, Douglas Procunier J (2006). Identification of single-nucleotide polymorphisms linked to resistance gene *Pc68* to crown rust in cultivated oat. Canadian journal of plant pathology 28: 214–222. 10.1080/07060660609507289

Cuddy W, Park R, Bariana H, Bansal U, Singh D, Roake J, Platz G (2016). Cereal rust report 2016. Plant Breeding Institute, University of Sydney.

Dodds PN (2023). From gene-for-gene to resistosomes: Flor’s Enduring Legacy. MPMI 36:461–446. 10.1094/MPMI-06-23-0081-HH

Fleischmann G, McKenzie RI (1968). Inheritance of crown rust resistance in *Avena Sterilis*. Crop Science 8:710–713.

Fleischmann G, McKenzie RIH, and Shipton WA (1971). Inheritance of crown rust resistance in *Avena sterilis* L. from Israel. Crop Science. 11:451–454

Flor HH (1971). Current status of gene-for-gene concept. Annu. Rev. Phytopathology. 9:275–296. 10.1146/annurev.py.09.090171.001423

Figueroa M, Dodds PN, Henningsen EC (2020). Evolution of virulence in rust fungi — multiple solutions to one problem. Curr. Opin. Plant Biol 56: 20–27. 10.1016/j.pbi.2020.02.007

Gnanesh BN, Mitchell Fetch J, Menzies JG et al (2013) Chromosome location and allele-specific PCR markers for marker-assisted selection of the oat crown rust resistance gene *Pc91*. Molecular Breeding 32:679–686. 10.1007/s11032-013-9900-6

Gnanesh BN, McCartney, CA, Eckstein PE, Mitchell Fetch JW, Menzies JG, Beattie AD (2015). Genetic analysis and molecular mapping of a seedling crown rust resistance gene in oat. TAG. Theoretical and applied genetics. Theor Appl Genet, 128(2), 247–258. 10.1007/s00122-014-2425-5

Haug-Baltzell A, Stephens SA, Davey S, Scheidegger CE, Lyons E (2017). SynMap2 and SynMap3D: web-based whole-genome synteny browsers. *Bioinformatics (Oxford*, England), 33(14), 2197–2198. 10.1093/bioinformatics/btx144

Henningsen EC, Lewis DC, Nguyen DT et al (2024). Virulence patterns of oat crown rust in Australia - season 2022. Plant Disease, 108:1959–1963. 10.1094/PDIS-09-23-1973-SC

Hewitt T, Henningsen EC, Pereira DA et al (2023). Genome-enabled analysis of population dynamics and virulence associated loci in the oat crown rust fungus *Puccinia coronata* f. sp. *avenae*. Molecular Plant-microbe Interactions. 10.1094/mpmi-09-23-0126-fi

Hoffman DL, Chong J, Jackson EW, Obert DE (2006). Characterization and mapping of a crown rust resistance gene complex *(Pc58)* in TAM O-301. Crop Science. 46:2630–2635. 10.2135/cropsci2006.01.0014

Kebede AZ, Friesen-Enns J, Gnanesh BN et al (2019). Mapping oat crown rust resistance gene *Pc45* confirms an Association with *PcKM*. G3 Genes|Genomes|Genetics 9:505–511. 10.1534/g3.118.200757

Klenová-Jiráková H, Leišová-Svobodová L, Hanzalová A, Kučera L (2010). Diversity of oat crown rust (*Puccinia coronata* f.sp. *avenae*) isolates detected by virulence and AFLP analyses. Plant Protect. Sci. 46:98–106. 10.17221/17/2009-PPS

Klos KE, Admassu-Yimer B, Babiker E et al (2017). Genome-wide association mapping of crown rust resistance in oat elite germplasm. The Plant Genome 10: 10.3835/plantgenome2016.10.0107

Kulcheski FR, Graichen FAS, Martinelli JA et al (2010). Molecular mapping of *Pc68*, a crown rust resistance gene in *Avena sativa*. Euphytica 175: 423–432. 10.1007/s10681-010-0198-8

Mammadov J, Aggarwal R, Buyyarapu R, Kumpatla S (2012). SNP markers and their impact on plant breeding. International Journal of Plant Genomics 2012:1–11. 10.1155/2012/728398

McCartney CA, Stonehouse RG, Rossnagel BG et al (2010). Mapping of the oat crown rust resistance gene *Pc91*. Theoretical and Applied Genetics 122:317–325. 10.1007/s00122-010-1448-9

McMullen MS, Doehlert DC, Miller JD (2005). Registration of “Morton” oat. Crop Science 45:1664–1665. 10.2135/cropsci2005.004

McNish I, Zimmer CM, Susko AQ et al (2020). Mapping crown rust resistance at multiple time points in elite oat germplasm. The Plant Genome 13: e20007 10.1002/tpg2.20007

Miller ME, Zhang Y, Omidvar V, Sperschneider J, Schwessinger B, Raley C, Palmer JM, Garnica D, Upadhyaya N, Rathjen J, Taylor JM, Park RF, Dodds PN, Hirsch CD, Kianian SF, Figueroa M (2018). *De Novo* Assembly and phasing of dikaryotic genomes from two isolates of *Puccinia coronata* f. sp. *avenae*, the causal agent of oat crown rust. mBio 9:10.1128/mbio.01650-17. 10.1128/mbio.01650-17

Miller ME, Nazareno ES, Rottschaefer SM, Riddle J, Dos Santos Pereira D, Li F, Nguyen-Phuc H, Henningsen EC, Persoons A, Saunders DGO, Stukenbrock E, Dodds PN, Kianian SF, Figueroa M (2020). Increased virulence of *Puccinia coronata* f. sp. *avenae* populations through allele frequency changes at multiple putative *Avr* loci. PLOS Genet. 16(12), e1009291. 10.1371/journal.pgen.1009291

Mitchell Fetch JW, Duguid SD, Brown PD et al (2007). Leggett oat. Canadian Journal of Plant Science 87:509–512. 10.4141/cjps06030

Moreau ELP, Riddle JM, Nazareno ES, Kianian SF (2024). Three decades of rust surveys in the United States reveal drastic virulence changes in oat crown rust. Plant disease, PDIS09231956RE. Advance online publication. 10.1094/PDIS-09-23-1956-RE

Mundt CC (2018). Pyramiding for resistance durability: Theory and practice. Phytopathology, 108:792–802. 10.1094/PHYTO-12-17-0426-RVW

Nazareno ES, Li F, Smith M et al (2017). *Puccinia coronata* f. sp. *avenae*: a threat to global oat production. Molecular Plant Pathology 19:1047–1060. 10.1111/mpp.12608

Nazareno ES, Fiedler JD, Miller ME et al (2022). A reference-anchored oat linkage map reveals quantitative trait loci conferring adult plant resistance to crown rust (*Puccinia coronata* f. sp. *avenae*). Theoretical and Applied Genetics 135:3307–3321. 10.1007/s00122-022-04128-6

Nguyen DT, Henningsen EC, Lewis D et al (2023). Genotypic and resistance profile analysis of two oat crown rust differential sets urges coordination and standardisation. Phytopathology. 114(6), 1356–1365. 10.1094/phyto-10-23-0353-r

Nguyen DT, Henningsen EC, Lewis D, Mago R, Sperschneider J, Stone E, Dodds PN, Figueroa M (2024). Characterisation of virulence of *Puccinia coronata* f. sp. *avenae* in Australia in the 2023 growing season. bioRxiv 2024.09.25.614061; doi: 10.1101/2024.09.25.614061

Park R, Wellings C, Bariana H, Bansai U (2009). Australia cereal cultivar pedigree and seedling rust genotype information. Cereal Rust Report 2009. Plant Breeding Institute, University of Sydney. https://www.sydney.edu.au/content/dam/corporate/documents/sydney-institute-of-agriculture/research/plant-breeding-and-production/cereal_rust_report_2009_vol_7_2.pdf

Park R (2013). New oat crown rust pathotype with virulence for *Pc91*. Cereal Rust Report 2013. Plant Breeding Institute, University of Sydney. https://www.sydney.edu.au/content/dam/corporate/documents/sydney-institute-of-agriculture/research/plant-breeding-and-production/cereal_rust_report_2013_vol_11_1.pdf

Park R, Chhetri M, Singh D (2023). Rust resistance genotypes and expected rust responses of Australian cereal varieties. Cereal Rust Report 2023. Plant Breeding Institute, University of Sydney. https://www.sydney.edu.au/content/dam/corporate/documents/faculty-of-science/research/life-and-environmental-sciences/rust-reports/cereal-rust-report-2023-vol-20-3.pdf

Periyannan S, Milne RJ, Figueroa M, Lagudah ES, Dodds PN. An overview of genetic rust resistance: From broad to specific mechanisms. PLoS Pathog. 2017 Jul 13;13(7):e1006380. 10.1371/journal.ppat.1006380

Rahman MZ, Hasan MT, Rahman J (2023). Kompetitive Allele-Specific PCR (KASP): An efficient high-throughput genotyping platform and its applications in crop variety development. In: Kumar, N. (eds) Molecular Marker Techniques. Springer, Singapore. 10.1007/978-981-99-1612-2_2

Satheeskumar S, Sharp PJ, Lagudah E, et al (2011). Genetic association of crown rust resistance gene *Pc68*, storage protein loci, and resistance gene analogues in oats. Genome 54:484–497. 10.1139/g11-014

Shi P, Shen X, Chen J-C et al (2023). KASP genotyping and semi-quantitation of G275E mutation in the *α6* subunit of *Thrips palmi*nAChR gene conferring spinetoram resistance. Pest Management Science 79:1777–1782. 10.1002/ps.7353

Simons MD (Marr Dixon) (1978). Oats: A standardized system of nomenclature for genes and chromosomes and catalog of genes governing characters. Washington, D.C: U.S. Dept. of Agriculture, Science and Education Administration. Print. https://books.google.com.au/books?id=iHyVsjyOiSEC

Simons MD (1985). Crown rust. Elsevier eBooks 131–172. 10.1016/b978-0-12-148402-6.50013-4

Sowa S, Paczos-Grzęda E (2020). Identification of molecular markers for the *Pc39* gene conferring resistance to crown rust in oat. Theoretical and Applied Genetics 133:1081–1094. 10.1007/s00122-020-03533-z

Strunk C, Kleinjan J, Caffe, M (2022, June 16). Crown Rust of Oats. SDSU Extension. https://extension.sdstate.edu/crown-rust-oats-0

van Niekerk B, Pretorius ZA, Boshoff WHP (2001). Pathogenic variability of *Puccinia coronata* f. sp. *avenae* and *P. graminis* f. sp. *avenae* on oat in South Africa. Plant disease 85, 1085–1090. 10.1094/PDIS.2001.85.10.1085

Wight CP, O’Donoughue LS, Chong J et al (2004). Discovery, localization, and sequence characterization of molecular markers for the crown rust resistance genes *Pc38*, *Pc39*, and *Pc48* in cultivated oat (*Avena sativa* L.). Molecular Breeding 14:349–361. 10.1007/s11032-005-0148-7

Wight CP, Blake VC, Jellen EN, Yao E, Sen TZ, Tinker NA (2024). One hundred years of comparative genetic and physical mapping in cultivated oat (*Avena sativa*). Crop & Pasture Science 75, CP23246. 10.1071/CP23246

Wong LSL, McKenzie RIH, Harder DE, Martens JW (1983). The inheritance of resistance *to Puccinia coron*ata and floret characters in *Avena sterilis*. Canadian Journal of Genetics and Cytology 25: 329–335. 10.1139/g83-052

Zhao J, Kebede AZ, Bekele WA et al (2020). Mapping of the oat crown rust resistance gene *Pc39* relative to single nucleotide polymorphism markers. Plant Disease. 104:1507–1513. 10.1094/pdis-09-19-2002-re

